# Bacterial Infection Remodels the DNA Methylation Landscape of Human Dendritic Cells

**DOI:** 10.1101/016022

**Authors:** Alain Pacis, Ludovic Tailleux, John Lambourne, Vania Yotova, Anne Dumaine, Anne Danckaert, Francesca Luca, Jean-Christophe Grenier, Kasper D Hansen, Brigitte Gicquel, Miao Yu, Athma Pai, Jenny Tung, Chuan He, Tomi Pastinen, Roger Pique-Regi, Yoav Gilad, Luis B Barreiro

## Abstract

DNA methylation is thought to be robust to environmental perturbations on a short time scale. Here, we challenge that view by demonstrating that the infection of human dendritic cells (DCs) with a pathogenic bacteria is associated with rapid changes in methylation at thousands of loci. Infection-induced changes in methylation occur primarily at distal enhancer elements, including those associated with the activation of key immune-transcription factors and genes involved in the crosstalk between DCs and adaptive immunity. Active demethylation is associated with extensive epigenetic remodeling and is strongly predictive of changes in the expression levels of nearby genes. Collectively, our observations show that rapid changes in methylation play a previously unappreciated role in regulating the transcriptional response of DCs to infection.

---

The first immune mechanisms recruited to defend against invading pathogens are those associated with innate immune cells, such as dendritic cells (DCs) or macrophages. Once they sense an intruder, these cells induce sophisticated transcriptional programs involving the regulation of thousands of genes, which are coordinated with the help of signal-dependent transcription factors, including NF-κB/Rel, AP-1, and interferon regulatory factors (IRFs)^1,2^. The regulation of this program is achieved through a series of epigenetic changes, which modulate the access of transcription factors to specific DNA regulatory elements^3^.

The most well-studied epigenetic responses to immune stimuli involve the post-translational modification of histone tails at promoter and enhancer regions^3,4^. Histone acetylation has been shown to be essential for the activation of many pro-inflammatory genes^5,6^, whereas increased activity of histone deacetylases is often associated with gene repression in the context of inflammation^7^. Moreover, recent studies suggest that the response of innate cells to different immune challenges can result in the appearance of histone marks associated with *de novo* enhancer elements (or latent enhancers)^8,9^, which have been postulated to contribute to a faster and stronger transcriptional response to a secondary stimulus^8^.

In contrast, we still know remarkably little about the role of other epigenetic changes in controlling responses to infection. DNA methylation has been particularly understudied, as a consequence of the belief that methylation marks are highly stable, and unlikely to respond to environmental perturbations on a short time scale. Recent work, however, suggests that DNA methylation patterns can rapidly change in response to certain environmental cues^10–13^, raising the possibility that rapid changes in DNA methylation might play a role in innate immune responses. Yet, to date, no studies have comprehensively investigated the contribution of rapid, active changes in methylation (in contrast to passive changes during cell replication) to the regulatory programs induced by innate immune cells in response to an infectious agent. More broadly, the few studies that demonstrate active regulation of DNA methylation in mammalian cells have been focused on a limited number of CpG sites and, surprisingly, the changes observed have been poorly predictive of changes in gene expression levels^10,12,13^. Here, we show that active changes in DNA methylation are pervasive in response to infection and, in contrast to previous reports; we found that such changes have a strong predictive impact on gene expression levels. Specifically, we report the first comprehensive epigenome and transcriptome of monocyte-derived DCs – professional antigen presenting cells that play a central role in bridging innate and adaptive immunity – before and after *in vitro* infection with live pathogenic bacteria. Our results thus provide unique insight into the role of active changes in methylation and their association with other epigenetic changes in the control of innate immune responses to infection.

## RESULTS

### MTB infection induces active changes in DNA methylation in human DCs

We infected monocyte-derived DCs from six healthy donors with a live virulent strain of *Mycobacterium tuberculosis* (MTB), the causative agent of tuberculosis (TB) in humans. Monocyte-derived DCs are ideally suited to study active changes in methylation given that they are post-mitotic and that they do not proliferate in response to infection^14,15^. At 18 hours after infection, we obtained paired data on single-base-pair resolution DNA methylation levels (using whole genome shotgun bisulfite sequencing: i.e., MethylC-Seq) and genome-wide gene expression data (using mRNA sequencing: i.e., mRNA-Seq) in non-infected and MTB-infected DCs. For MethylC-Seq data, we generated 8.6 billion single-end reads (mean of 648 ± 110 SD million reads per sample; **Supplementary Table 1**) resulting in an average coverage per CpG site of ~9X for each sample. We detected an average of 24 million CpG sites in each sample, corresponding to over 80% of CpG sites in the human genome. Genome-wide methylation data between biological replicates was strongly correlated attesting for the high quality of the data (**Supplementary Fig. 1**; mean r across all samples = 0.86).

As expected for mammalian cells, CpG sites were ubiquitously methylated throughout the genome except near transcription start sites (TSSs) and in CpG islands (**Supplementary Fig. 2a,b**). We found a significant negative correlation between gene expression levels and methylation levels around TSSs (r = −0.39; *P* < 1 × 10^−16^; **Supplementary Fig. 2c,d**), highlighting the well-established role of proximal methylation in the stable silencing of gene expression. Principal component analysis of our data along with MethylC-Seq data from 21 other purified cell types and tissues revealed that the DC methylome is closely related to that of other blood-derived cells, particularly cells that share a common myeloid progenitor with DCs, such as neutrophils (**Supplementary Fig. 2e**).

We next assessed the occurrence and the extent to which the response of DCs to a bacterial infection is accompanied by active changes in DNA methylation, using the BSmooth algorithm^16^. We defined MTB-induced differentially methylated regions (MTB-DMRs) as regions of 3 or more consecutive CpG sites exhibiting a significant difference in methylation between the two groups (*P* < 0.01) and an absolute mean methylation difference above 0.1^17^. Using these criteria, we identified 3,271 MTB-DMRs, within which 1,557 corresponded to hypermethylated regions (i.e., regions in which methylation levels increased upon infection; mean length = 134 bp) and 1,714 to hypomethylated regions (i.e., regions in which methylation levels decreased upon infection; mean length = 184 bp) (**Fig. 1a** and **Supplementary Table 2**). Interestingly, we found that less than 7% of MTB-DMRs overlapped with a putative promoter (**Fig. 1b**) and that the vast majority of these sites were located distal to TSSs (median distance of ~35 kb from the nearest TSS; **Fig. 1c**). MTB-DMRs occur in genomic regions that show increased levels of evolutionary conservation (**Fig. 1d**), a finding supporting that they are functionally important. Moreover, gene ontology analysis revealed that MTB-DMRs are significantly enriched (false discovery rate (FDR) < 0.05) near genes known to play a key role in the regulation of immune processes, including the regulation of transcription, signal transduction, and, importantly, cell apoptosis – a critical mechanism assisting DC cross-presentation to adaptive immune cells (**Fig. 1e**). Notably, hypomethylated regions show a stronger enrichment for genes involved in DC-T cell crosstalk, suggesting that active demethylation plays a particularly important role in dictating the nature and magnitude of the T cell response to infectious agents (**Fig. 1e**).

**Figure 1.**
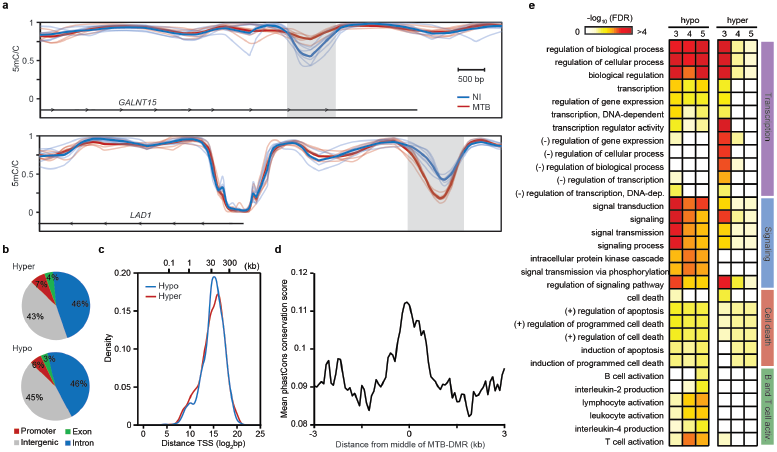
MTB-induced changes in methylation in human DCs. (**a**) Examples of regions showing active changes in methylation in response to MTB infection. The top panel shows an example of a hypermethylated region and the bottom an example of a hypomethylated region. The plot shows smoothed methylation values (y-axis) for six non-infected (blue) and six MTB-infected samples (red). Gray shadings highlight MTB-DMRs and thick blue and red lines show average methylation levels for non-infected and infected cells, respectively. (**b**) Pie charts showing the distribution of hyper- and hypomethylated regions in different genomic regions. Each MTB-DMR is counted only once: the overlap of a genomic region excludes all previously overlapped MTB-DMRs clockwise from promoters (TSS ± 500 bp; red) (**c**) Distribution of distances of MTB-DMRs to the nearest TSS. (**d**) Average conservation score (phastCons ^46^ score per 50 bp) around the center of MTB-DMRs not associated with promoter regions (i.e, > 3kb away from any known TSS). (**e**) Representative gene ontology (GO) terms enriched among genes associated to hyper- and hypomethylated regions. To demonstrate that the enriched biological processes are largely robust to the cutoff used to define MTB-DMRs we show the enrichment results when requiring 3+ (the one used on the main text), 4+ or 5+ consecutive differently methylated CpG sites (*P*<0.01) to call an MTB-DMR.

### Active Changes in Methylation Occur in Regions Enriched for 5-hydroxymethylcytosine

The TET family proteins catalyze the conversion of methylated cytosine (5mC) to 5-hydroxy-methylcytosine (5hmC), and are thus key players in the process of active demethylation. To evaluate if 5hmC were dynamically changing in response to MTB infection (as expected if 5mC sites must pass through the 5hmC state before demethylation), we generated single-base pair resolution maps of 5hmC across the genome using Tet-assisted bisulfite sequencing (TAB-Seq)^18^ in one additional donor. As previously described for other cell populations^19,20^, we found markedly higher levels of 5hmC in gene bodies of highly expressed genes, consistent with a role for 5hmC in maintaining and/or promoting gene expression (**Fig. 2a**)^21,22^.

**Figure 2.**
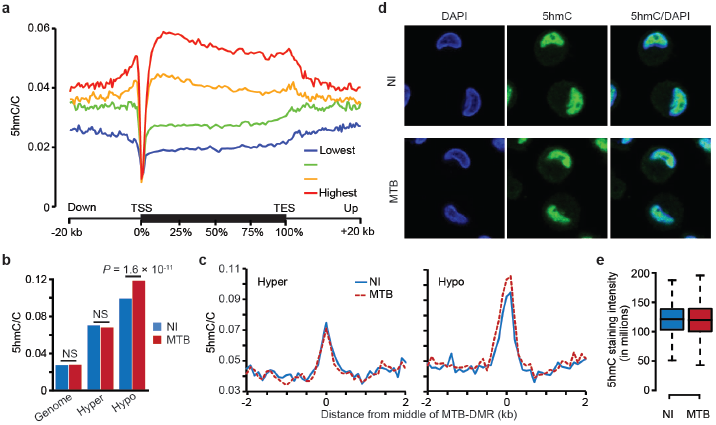
5hmC is enriched in MTB-DMRs prior to infection. (**a**) Metagene profiles of 5hmC levels relative to Ensembl transcripts expressed at different levels in human DCs. We grouped genes in four quantiles based on their expression levels on non-infected DCs. (**b**) Barplots showing mean 5hmC/C ratios within hyper- and hypomethylated regions, before (blue) and after infection (red). (**c**) Composite plots of patterns of 5hmC before and after MTB infection ±3 kb around the midpoint of hyper- and hypomethylated regions. (**d**) 5hmC staining in non-infected (top panel) and MTB-infected DCs (bottom panel). 5hmC levels are given by the levels of Alexa 488 (green: middle panel) and cells counterstained with DAPI to localize the nucleus (first panel). (**e**) Boxplots showing the distribution of 5hmC staining intensity. No significant differences were observed between the two groups.

Next, we evaluated if 5hmC marks were enriched within MTB-DMRs. We found that regions in which methylation actively changed in response to infection were already associated with significantly higher levels of 5hmC prior to infection (2.6- and 3.6-fold enrichments for hyper-and hypomethylated regions; Wilcoxon test; *P* < 1 × 10^−16^, respectively). Upon infection, hypomethylated regions show increased levels of 5hmC (Wilcoxon test; *P* = 1.57 × 10^−11^; **Fig. 2b,c**), which suggests that 5hmC plays an important role in the cascade of events leading to active demethylation. The increase in 5hmC appears to be specific to hypomethylated regions: no enrichment was observed in hypermethylated regions or genome-wide, a result supported by quantitative immunocytochemistry data (**Fig. 2d,e**). The striking enrichment of 5hmC within MTB-DMRs prior to infection strongly suggests that, in addition to its role as a transitory demethylation intermediate, 5hmC might also serve additional regulatory functions including the coordination of the gene expression program induced in response to a microbial stimulus.

### MTB-DMRs overlap with enhancer elements that gain activation marks upon infection

Given that MTB-DMRs are primarily found distal to TSSs, we predicted that MTB-DMRs would overlap with enhancer regions. To test this hypothesis and evaluate how the chromatin states associated with MTB-DMRs might dynamically change in response to infection, we collected ChIP-Seq data for six histone marks (H3K4me1, H3K4me3, H3K27ac, H3K27me3, H3K36me3 and H3K9me3) in non-infected and infected DCs (**Supplementary Table 1**) from two additional donors. Using these data, we generated genome-wide, gene regulatory annotation maps for non-infected and MTB-infected DCs using the ChromHMM chromatin segmentation program (**Fig. 3a** and **Supplementary Fig. 3**)^23^. We found that 37% of MTB-DMRs overlapped with a ChromHMM-annotated enhancer region (defined by the presence of H3K4me1) already present in non-infected DCs, a 6.7-fold enrichment compared to genome-wide expectations (*χ2*-test; *P* < 1 × 10^−16^; **Fig. 3b,c**, and **Supplementary Table 2**). Similar enrichments (6.7-fold; *P* < 1 × 10^−16^) were observed when defining chromatin states in MTB-infected DCs. Given the high-resolution of our histone maps, we could further distinguish between active and inactive/poised enhancer elements based on the presence or absence of the H3K27ac mark, respectively, in addition to H3K4me1^24–26^. Overall, we found that MTB infection leads to a significant increase of active enhancer elements (and decrease of inactive/poised enhancers) colocalizing with MTB-DMRs (**Fig. 3b,c**).

**Figure 3.**
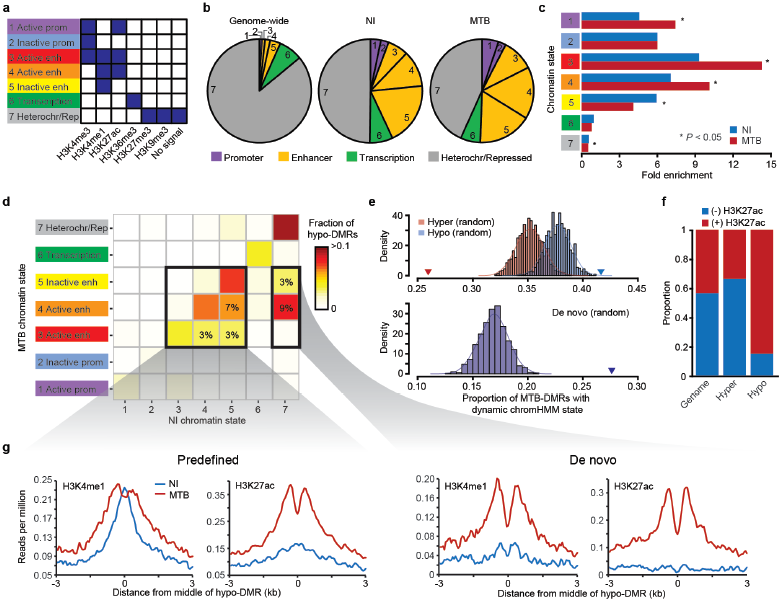
MTB-DMRs overlap with enhancer elements that become active upon infection in hypomethylated regions. (**a**) Combination of histone patterns used to define the 7 chromatin states. The precise relative contribution of each chromatin mark to each of the chromHMM-defined states can be found in **Supplementary Fig. 3**. Note that state 7 was defined by either no signal *or* the presence of either H3K27me3/H3K9me3. (**b**) Pie charts showing the distribution of chromatin state annotations genome-wide (on non-infected cells) and within all MTB-DMRs in either non-infected (blue) or MTB-infected cells. The chromatin state codes are as defined in (**a**). (**c**) Fold enrichments of the different chromatin states within MTB-DMRs as compared to genome-wide expectations in non-infected (blue) and MTB-infected cells (red). (**d**) Heatmap of the proportion of hypomethylated regions by chromatin transition state. The x-axis represents the chromatin states defined in non-infected DCs and the y-axis the chromatin state of the same region in MTB-infected DCs. The two inner boxes indicate two subgroups of hypomethylated regions, predefined enhancer (detectable enhancer in non-infected DCs) and *de novo* enhancers (detectable enhancer only in MTB-infected DCs). The numbers inside the cells refer to the proportion of hypomethylated regions that undergo each of the highlighted transitions. (**e**) Top panel: Histogram showing the observed proportion of regions that change chromatin state after infection (any transition) when sampling 1000 random sets of regions matched to the chromatin states found in non-infected samples within hypomethylated regions (blue) and hypermethylated regions (red). Each random set contains the same number of hypomethylated and hypermethylated regions as those identified in the true data. The blue and red triangles represent the observed proportion of regions that changed chromatin state in response to MTB infection in hypomethylated and hypermethylated regions, respectively. Bottom Panel: Same as above but focusing on regions of the genome labeled as heterochromatin/repressed before infection (state 7) that gain *de novo* enhancer marks upon MTB infection (states 3, 4, or 5). The blue triangle represents the proportion observed within the true set of hypomethylated regions. (**f**) Barplots showing the proportion of hyper- and hypomethylated regions that overlap with enhancers and show dynamic changes in chromatin state, as defined by the gain or loss of H3K27ac mark. (**g**) Composite plots of patterns of H3K4me1 and H3K27ac ChIP-Seq signals ±3 kb around the midpoints of hypomethylated regions (x-axis) overlapping with predefined (right) and *de novo* (left) enhancers.

Previous studies have shown that enhancer activation and inactivation are associated with local CpG hypomethylation and hypermethylation, respectively^20^. We therefore extended our analysis by examining chromatin transition states at hyper- and hypomethylated regions. In hypermethylated regions, we found no evidence for an enrichment in the proportion of regions whose chromatin state annotation changed in response to infection, compared to the genomewide background (**Fig. 3e** and **Supplementary Fig. 4**). Conversely, 42% of hypomethylated regions occurred in regions that exhibited infection-dependent changes in chromatin state, a significantly higher proportion than expected compared to the rest of the genome (*P*_resampling_ < 0.001; **Fig. 3e**). The chromatin state transitions observed within hypomethylated regions were primarily explained by the acquisition of histone activating marks (e.g., H3K27ac) in MTB-infected cells. For example, among hypomethylated regions that overlapped with predefined enhancers (i.e., enhancers observable in non-infected cells), 85% of those that exhibit a change in chromatin state gained an activation mark (H3K27ac or H3K27ac+H3K4me3; **Fig. 3f,g** and **Supplementary Fig. 5**). This proportion was markedly larger than that observed in hypermethylated regions (34%) or genome-wide (44%) (*χ2*-test; *P* < 8.3 × 10^−5^; **Fig. 3f**). Notably, we also found a large number of hypomethylated regions (n = 218; 12.7% of all hypomethylated DMRs) that overlapped with heterochromatin/repressed regions before infection but gained *de novo* enhancer marks upon MTB infection (H3K4me1 (+ H3K27ac + H3K4me3)). The number of *de novo* enhancers we observed among hypomethylated DMRs was significantly higher than expected by chance (*P*_resampling_ < 0.001; **Fig. 3d,e,g** and **Supplementary Fig. 5**). The identification of enhancers only present in infected DCs resembles recent findings in mouse macrophages showing that in response to different immune stimuli they can gain *de novo* putative enhancer regions that were absent in naive cells^8,9^.

Interestingly, MTB-induced activation or *de novo* gain of enhancer elements at hypomethylated regions is associated with the induction of putative enhancer RNAs (eRNAs)^27^ in these intergenic regions (as measured by whole-transcriptome RNA-seq) as well as with increased levels of histone marks associated with transcriptional activity (**Supplementary Fig. 6**). Moreover, changes in eRNA levels in response to MTB infection show a striking positive correlation with changes in gene expression levels of nearby genes (r = 0.49, *P* = 7.6 × 10^−13^; **Supplementary Fig. 6**), in support to a mechanistic link between demethylation, eRNA production and the regulation of proximal protein-coding genes^28^.

### MTB-DMRs are bound by signal-dependent transcription factors

We next asked if MTB-infection was associated with changes in the levels of chromatin accessibility in MTB-DMRs. We mapped regions of open chromatin in non-infected and infected DCs based on genome-wide sequencing of regions showing high transposase (Tn5) sensitivity (using ATAC-Seq in one additional donor)^29^. Overall, we observed that MTB-DMRs colocalize with regions of open chromatin, which further reinforces the regulatory potential of these regions (**Fig. 4a**). Interestingly, we found that the response to MTB-infection was accompanied by a striking increase in Tn5 sensitivity levels in hypomethylated regions, which indicates that the chromatin in these regions became more accessible after infection (**Fig. 4a**). This observation is commensurate with our data showing the acquisition of active histone marks in these regions, and further supports the idea that hypomethylated regions frequently reflect the presence of regulatory elements that become more active in response to infection.

**Figure 4.**
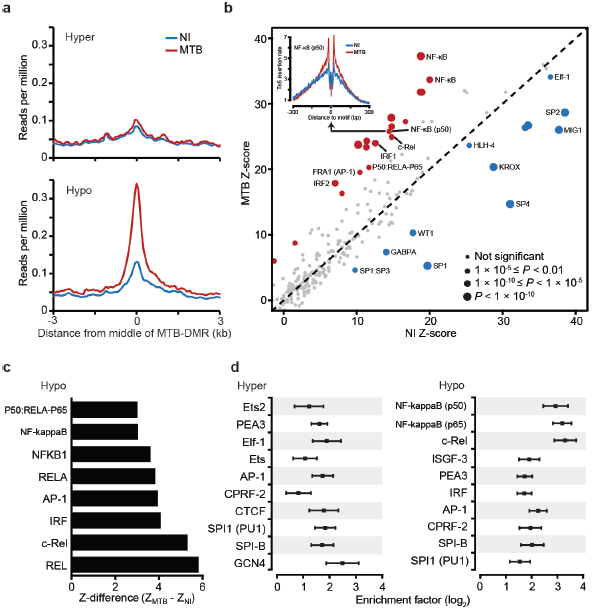
MTB-DMRs are bound by signal-dependent transcription factors. (**a**) Tn5-accessibility profiles before and after MTB infection, ±3 kb around the midpoints of hypermethylated regions (top panel) and hypomethylated regions (bottom panel). (**b**) Scatterplot comparing transcription factor occupancy score predictions between non-infected (y-axis) and MTB infected DCs (y-axis). The size of the dots is proportional to the level of statistical significance supporting differential binding in response to MTB infection. Red dots represent TFs that show evidence for increased binding after MTB infection and blue dots represent TFs that show evidence for decreased binding after infection. The inset on the top right corner shows the genome-wide footprint for NF-κB (p50) (motif ID: M00051) in non-infected (blue) and MTB-infected DCs (red). In this example, the footprint in MTB-infected DCs is clearly stronger, which supports increased binding of NF-κB (p50) genome-wide upon MTB infection. (**c**) TF motifs that show significantly increased binding in hypomethylated regions after MTB infection. (**d**) TFs for which the number of well-supported footprints (posterior Pr > 0.95) within hyper- and hypomethylated regions were enriched relative to the genomic background, in MTB infected DCs. The enrichment factors are shown in the x-axis in a log2 scale. The bars around the estimated enrichments reflect the 95^th^ confidence intervals around the estimates. For visualization purposes we only show the top 10 most significantly enriched TF footprints. A complete list of all TFs for which footprints are enriched within MTB-DMRs can be found in **Supplementary Table 3**.

An attractive feature of ATAC-Seq data is the ability to identify motif instances occupied by transcription factors (TF) within regions of open chromatin^29,30^. We did so by using a modified version of the Centipede algorithm^31^ specifically devised to test for aggregate differential binding of TFs between two experimental conditions. This method, which we coined CentiDual, compares the intensity of the Tn5 sensitivity-based footprint across all matches to a given motif in the genome, between non-infected and infected samples (see Online Methods for details on the statistical model). We found compelling evidence for measurable, genome-wide transcription factor activity (i.e., binding to the genome; Bonferroni-corrected *P* < 0.05) in either non-infected or infected DCs for 264 TF binding motifs, representing over 200 unique transcription factors (some TFs can bind different motifs; **Supplementary Table 3**). Of these TF binding motifs, we found 56 that were differentially bound between non-infected and infected DCs (Bonferroni-corrected *P* < 0.05; 26 show increased binding and 29 show decreased binding; **Fig. 4b**). Among TF binding motifs showing increased genome-wide binding after infection, we found several NF-κB and IRF family members (**Fig. 4b** and **Supplementary Table 3**), both of which play a primary role in the regulation of inflammatory signals in response to infection^2^. Interestingly, several CTCF motifs showed significantly decreased binding in infected DCs (Bonferroni-corrected *P* < 1.85 × 10^−14^, **Supplementary Table 3**). CTCF is a well-established transcriptional insulator^32^, raising the possibility that the release of CTCF in response to infection might represent an important mechanism for the regulation of efficient immune responses.

We next used CentiDual to test for differential binding within MTB-DMRs. We found no significant evidence of differential binding for any of the 264 TF binding motifs tested within hypermethylated regions. In contrast, within hypomethylated regions we found increased binding (FDR < 0.1) at 8 TF binding motifs after infection (**Supplementary Table 3**). Strikingly, all of these motifs were associated with immune-induced TFs from the NF-κB (e.g., REL; FDR = 1.57 × 10^−6^), AP-1 (FDR = 4.9 × 10^−3^), or IRF (FDR = 3.97 × 10^−3^) families (**Fig. 4c**), which demonstrates that hypomethyated regions correspond to places where immune-activated TFs are being recruited. In accordance, we found that, in infected DCs, NF-κB, AP-1, and IRFs were all significantly enriched within MTB-DMRs (up to 16-fold), particularly within hypomethylated regions (**Fig. 4d**). Indeed, in MTB infected DCs, over 50% of the hypomethylated regions were bound by at least one of these signal-dependent TFs, which corresponds to an 3.8-fold increase relative to expectation (based on sampling random regions of the genome matched for length and GC content; **Supplementary Fig. 7**; *χ2*-test; *P* = 3.94 × 10^−63^).

### MTB-DMRs are associated with genes differently expressed in response to MTB infection

Finally, we asked if genes associated with MTB-DMRs were more likely to change expression levels in response to infection. We classified 1,665 and 1,740 genes as significantly up- or down-regulated post-infection, respectively (FDR < 1 × 10^−4^ & |log2 fold-change| > 1; **Supplementary Table 4**). We next tested whether genes located nearby MTB-DMRs were more likely to be differentially expressed upon MTB infection relative to all genes in the genome. To do so, we first associated each MTB-DMR with a unique gene using the following criteria: if an MTB-DMR was located within a gene body, the MTB-DMR was assigned to that gene; otherwise, we assigned each MTB-DMR to the gene with the TSS closest to the midpoint of the MTB. Then, we tested for an enrichment of differentially expressed (DE) genes among five classes of genes: (*i*) “DMR-genes” corresponding to all genes associated with MTB-DMRs (n = 2,160); (*ii*) “hyper-DMR-genes” corresponding to the set of genes associated with hypermethylated regions (n = 1,137); (*iii*) “hypo-DMR-genes” corresponding to the set of genes associated with hypomethylated regions (n = 1,201); (*iv*) “predefined-DMR-genes” corresponding to the set of genes in hypomethylated regions that overlapped with predefined enhancer elements (n = 470, a subset of class iii), and (*v*) “*de novo*-DMR-genes” corresponding to the set of genes in hypomethylated regions that overlapped with *de novo* enhancer elements (n = 161, also subset of class *iii*).

We found that DMR-genes (class *i*) were significantly enriched among DE genes (1.4-fold, *χ2*-test; *P* = 1.27 × 10^−11;^ **Fig. 5a,b**) as compared to all genes in the genome. This enrichment was noticeably stronger for predefined-DMR-genes (class *iv*; 1.9-fold, *P* = 3.1 × 10^−11^) and even more so for *de novo*-DMR-genes (class *v*; 2.3-fold, *P* = 1.1 × 10^−9^). Indeed, among *de novo*-DMR-genes, 49% were DE, even at very stringent cutoffs we used to define DE genes (**Fig. 5a,b**). Interestingly, among DE genes associated with hypomethylated regions, 73% were up-regulated after MTB infection – substantially more than the 51% of up-regulated genes observed genomewide (*χ2*-test; *P* = 1.41 × 10^−73^, **Fig. 5c**). This observation was even more pronounced when focusing specifically on predefined-DMR-genes (class *iv*) and *de novo*-DMR-genes (class *v*), for which 78% (*P* = 5.46 × 10^−34^) and 92% (*P* = 1.39 × 10^−19)^, respectively, were associated with increased expression levels in response to infection (**Fig. 5c**). Interestingly, the biological functions linked to predefined enhancer-associated genes differed from those linked to *de novo* enhancer-associated genes, even though both types of enhancers occurred in hypomethylated DMRs. Specifically, DE genes associated with predefined enhancer elements that loose methylation in response to infection were mainly enriched for signal transduction processes (FDR < 0.01). This set of genes included virtually all the key “master-regulators” of innate immune responses, such as *NFKB1*, *NFKB1A*, *IRF2*, and *IRF4* (**Fig. 5d**). In contrast, DE genes associated with *de novo* enhancers appear to be more directly involved in processes related to the ability of DCs to activate B and T cells (FDR < 0.01; e.g., *CD83*) and the regulation of cell death (FDR < 0.01; e.g., *BCL2*) (**Supplementary Table 5**).

**Figure 5.**
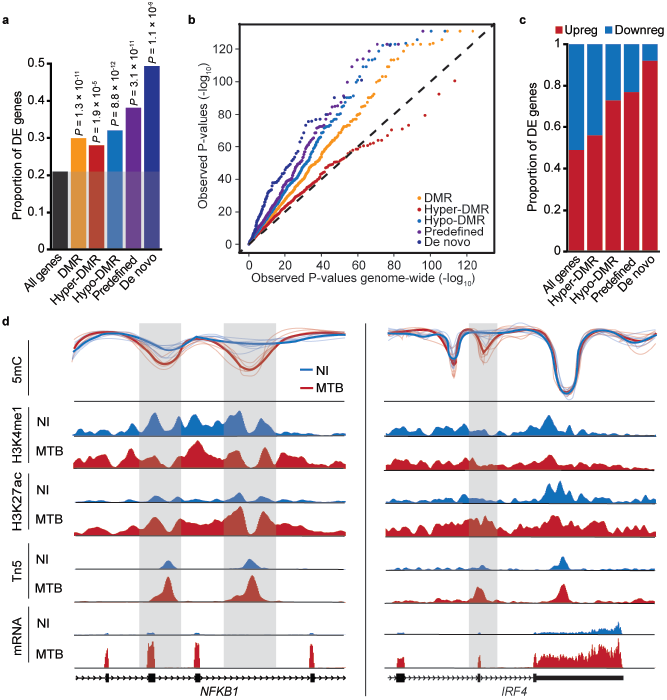
Differential methylation is coupled to differential gene expression. (**a**) Proportion of differentially expressed genes (y-axis) observed among all tested genes and among genes associated with different subgroups of MTB-DMRs. (**b**) QQ-plot showing that genes in the vicinity of MTB-DMRs (and particularly hypomethylated regions) show stronger statistical evidence for being differently expressed in response to MTB infection (*P* values on y-axis) compared to all genes tested (*P* values on x-axis). (**c**) Proportion of up- and down-regulated genes among DE genes associated with the different subgroups of MTB-DMRs. (**d**) Examples of genes encoding for two key transcription factors, *NFKB1* (left panel) and *IRF4* (right panel) that are strongly upregulated in response to MTB infection and for which we identified one or more hypomethylated regions (gray shading) that overlap with putative enhancer elements. Normalized read counts for the indicated features are shown for non-infected (blue tracks) and infected conditions (red tracks).

## DISCUSSION

The possibility that active changes in methylation, particularly demethylation, can occur in mammals has been a matter of debate for decades^33,34^. Here, we provide with compelling evidence that the response of human DCs to MTB infection is accompanied by widespread, rapid changes in DNA methylation that involve both active demethylation and *de novo* methylation. Several mechanisms could account for the observed active changes in methylation. *De novo* methylation is likely explained by the action of DNA methyltransferases (DNMT), namely DNMT3^35^. For demethylation many possible mechanisms can potentially be invoked^36^; however, the observation that hypomethylated regions show increased levels of 5hmc in response to MTB infection strongly suggests that the family of TET proteins (TET1-3) are involved in this process. This possibility is further supported by recent studies showing that TET2 was required for active DNA demethylation in human monocytes^13^ and during brain development^20^.

By integrating our methylation maps with ChIP-Seq data on six histone marks we show that active methylation changes occur almost exclusively at distal regulatory elements, namely enhancers. This observation, which is robust to the cutoffs used to call MTB-DMRs (**Supplementary Fig. 8**), parallels what has been previously described in differentiating cells and during developmental processes^37–39^ despite the fact that the mechanisms controlling active and passive changes in methylation are markedly different^36^. In stark contrast to previous studies that also reported active changes in methylation, for example, in response to neuronal activation^10^ or during the process of differentiation of human monocytes into macrophages or dendritic cells^13^, we found a strong association between DMRs and changes in gene expression of nearby genes. The apparent discrepancy between our results and those previously reported is probably explained by the fact that past studies have only investigated active methylation changes in promoter regions – that we show to be infrequent – or only on a small subset of all CpG sites in the (mouse) genome (~1%). Moreover, we decided to focus on differently methylated regions (3 or more consecutive differently methylated CpGs) instead of methylation changes at individual CpG sites^10,13^, which is likely to enrich for DMRs that are more directly linked to changes in gene expression. In support to that hypothesis, we found that the enrichment for DE genes become stronger as we focus on DMRs with a larger number of differently methylated CpG sites (**Supplementary Fig. 8**). More broadly, our results highlight the key importance of using single-base resolution map of the DNA methylome in order to fully capture the relationship between changes in methylation and changes in gene expression.

Interestingly, we found that the loss of methylation at enhancer elements is often associated with the gain of histone activation marks and the recruitment of immune-activated TFs (e.g., NF-κB and IRFs) to these regions in response to infection. The recruitment of NF-κB and other master regulators to hypomethylated regions is likely associated with the opening of the chromatin in these regions, although it remains unclear whether the chromatin opens to allow the binding of these TFs (i.e., prior to binding) or if the observed increase in chromatin accessibility is a consequence of the binding itself. It also remains to be further explored whether the observed changes in methylation are required to allow TF binding or if it is the binding of TFs to these regions that leads to the loss of methylation, as previously proposed in other cellular contexts^39,40^. Our analyses do indicate, however, that even if TF binding is the instigating factor to the observed changes in methylation, binding alone is not sufficient given that the vast majority (>99%) of binding events induced by infection occur at regions that do not change methylation (**Supplementary Fig. 9**).

There is mounting evidence that after a first encounter with a pathogen or other immune stimulus innate immune cells keep such attacks “in memory”, leading to faster and stronger gene transcriptional responses upon restimulation and increased resistance to secondary infection. This process, termed trained immunity^4,41,42^, has been attributed to epigenetic reprograming at the level of histone H3 methylation based on the observation that distal regulatory elements that gain *de novo* H3K4me1 (i.e., *de novo* enhancer marks) in response to immune activation generally do not lose this mark after the stimulation has ceased^8^. Although epigenetic programming through histone modifications might be an important factor in trained immunity, our results raise the possibility that changes in DNA methylation might also contribute to short-term memory in innate immune cells. Indeed, changes in DNA methylation might be ideally suited as a mechanism of epigenetic memory since these changes are expected to be thermodynamically more stable and longer lasting than changes in histone marks. Based on our current data, it is hard to assess the degree to which MTB-induced changes in DNA methylation remained stable over time. However, many of the genes associated with DMRs are well-recognized as early response genes (e.g., *NFKB1*, *IRF2*, *IRF4*, *IRF8* are upregulated minutes after an immune challenge^43,44^ or genes that only respond to MTB infection at early time points (e.g., *DUSP2, MAP2K3, MAP3K4*)^45^. This observation strongly suggests that some of the changes in methylation are likely to have occurred in the first hours after infection and have remained stable over time. Moreover, we show that the gain of *de novo* enhancers – assumed to account for trained immunity – often occurs concomitantly with the loss of DNA methylation in the same regions. Our results thus raise the possibility that trained immunity might not only be due to post-transcriptional changes in histone marks but also, and possibly primarily, due to changes in DNA methylation.

Collectively, our results suggest that active and rapid changes in methylation might play a previously unappreciated and critical role in the transcriptional regulation of an efficient immune response to infection. They also highlight the key importance of multiple and coordinated epigenetic changes in enhancer regions at dictating the regulatory programs induced in response to a live pathogenic bacteria.

## METHODS

Methods and any associated references are available in the online version of the paper.

*Note: Any Supplementary Information and Source Data files are available in the online version of the paper*.

## ACKNOWLEDGMENTS

We thank B. Jabri, V. Abadie and J.F. Brinkworth for helpful discussions and comments on the manuscript; K. Michelini and C. Chavarria for technical assistance running the sequencer; G. Stewart for a gift of the MTB strain used in this study; and P. Roux for advice on the confocal microscopy. This study was funded by National Institutes of Health Grant AI087658 (to Y.G. and L.T.), by grants from the Canadian Institutes of Health Research (301538 and 232519), the Human Frontiers Science Program (CDA-00025/2012) and the Canada Research Chairs Program (950-228993) (to L.B.B.), by the Canadian Institutes of Health Research/Canadian Epigenetics, Environment and Health Research Consortium (to T.P.) and by the NIH grant HG006827 (to C.H.). A.P. was supported by a fellowship from the Réseau de Médecine Génétique Appliquée (RMGA).

## AUTHOR CONTRIBUTIONS

A.P. performed data analysis and drafted the manuscript. L.T. performed the infection experiments with MTB and the quantitative immunocytochemistry with help from A. Danckaert. J.L. and T.P. performed ChIP-seq experiments for histone marks. V.Y. constructed bisulfite-treated DNA libraries and RNA-seq libraries. A.Dumaine. A.P. implemented the ATAC-seq protocol and generated the ATAC-seq libraries with help from F.L. developed J-C.G. provided assistance in the bioinformatics analysis. K.H. provided assistance with the identification of MTB-DMRs. M.Y. and C.H. performed the TAB-seq experiments. B.G. provide reagents for the infection experiments. J.T. contributed to the analysis and the writing of the manuscript. R.P-R. contributed new analytical tools for the fooptrinting analysis. Y.G. designed the study and provided with reagents and analytical tools. L.B.B. designed and supervised the study and primarily wrote the manuscript with contributions from all authors.

## ONLINE METHODS

**Sample collection.** Blood samples were obtained from the Indiana Blood Center. A signed written consent was obtained from all of the participants and the project was approved by the ethics committee at the CHU Sainte-Justine (protocol #4023). All individuals recruited in this study were healthy Caucasian males between the ages of 21 and 55 y old. We decided to only focus on males to limit the variation in methylation estimates due to sex-specific differences. Only individuals self-reported as currently healthy, not under medication, and with no history of chronic diseases were included in the study. In addition, each donor’s blood was tested for standard blood-borne pathogens, and only samples negative for all of the pathogens tested were included.

***Mycobacterium tuberculosis* preparation.** We infected dendritic cells (DCs) with a *Mycobacterium tuberculosis* (MTB) strain expressing green-fluorescent protein (H37Rv)^47,48^. Importantly, previous studies have shown that the presence of GFP in this strain does not alter the growth rate or the virulence of the bacilli under axenic conditions, relative to wild-type MTB. *M. tuberculosis* H37Rv was grown from a frozen stock to midlog phase in 7H9 medium (BD) supplemented with albumin-dextrose-catalase (ADC; Difco)^48,49^.

**Isolation and infection of DCs.** Peripheral blood mononuclear cells (PBMCs) were isolated from buffy coats by Ficoll-Paque centrifugation. Blood monocytes were then purified from PBMCs by positive selection with magnetic CD14 MicroBeads (Miltenyi Biotech). Pure monocytes were cultured for 5 days in RPMI 1640 (Invitrogen) supplemented with 10% heat-inactivated FCS (Dutscher), L-glutamine (Invitrogen), GM-CSF (20 ng/mL; Immunotools), and IL-4 (20 ng/mL; Immunotools). Cell cultures were fed every 2 days with complete medium supplemented with the cytokines previously mentioned. Before infection, we systematically checked the differentiation/activation status of the monocyte-derived DCs by flow cytometry, using antibodies against CD1a, CD14, CD83, and HLA-DR. Only samples presenting the expected phenotype for non-activated DCs – CD1a+, CD14-, CD83-, and HLA-DRlow – were used in downstream experiments. The resulting monocyte-derived DCs were then infected with MTB for 18 h at a multiplicity of infection of 1-to-1, as previously described^47^.

For biosecurity reasons the ChIP-Seq and ATAC-Seq experiments were performed using heat-killed bacteria instead of live MTB. In order to evaluate the extent to which using heat-killed bacteria could result in a different transcriptional response to that induced by live MTB, we used the Illumina HumanHT-12 v4 Expression BeadChip array to compare the genome-wide transcriptional responses observed in DCs in response to live MTB to those observed when DCs from the same donors were exposed to different amounts of heat-killed MTB bacteria. Low-level microarray processing including normalization of the data and variance stabilizing transformation were performed as previously described^47^. We found that using the equivalent of 5 heat-killed bacteria to 1 DC leads to virtually the same transcriptional response at 18 hours to that observed with live MTB (r = 0.91; **Supplementary Fig. 10**).

**DNA and RNA Extractions.** DNA from infected and non-infected DCs was extracted using the PureGene DNA extraction kit (Gentra Systems). Total RNA was extracted from the same samples using the miRNeasy kit (Qiagen). RNA quantity was evaluated spectrophotometrically, and the quality was assessed with the Agilent 2100 Bioanalyzer (Agilent Technologies). Only samples with no evidence for RNA degradation (RNA integrity number > 8) were kept for further experiments.

**MethylC-Seq library preparation and sequencing.** DNA from infected and non-infected DCs (6ug) was spiked with 30ng of unmethylated cl857 *Sam7* Lambda DNA (Promega, Madison, WI) and further sonicated to an average length of ~100bp using a Covaris ultrasonicator under the following settings for 16 cycles: Duty cycle: 10%; Intensity: 5; Cycles/burst: 100. The sonicated product was further subjected to repair of 3’ and 5’ ends followed by the addition of a non-templated dA-tail before ligation to cytosine-methylated adapters provided by Illumina (Illumina, San Diego, CA) as per manufacturerʼs instructions for genomic DNA library construction. Adapter-ligated DNA of 100-200 bp was isolated by 2% agarose gel electrophoresis, and sodium bisulfite conversion was performed on the resulting sample using the MethylCode™ Bisulfite Conversion Kit (Invitrogen) as per manufacturer’s instructions. Half of the bisulfite-converted, adapter-ligated DNA molecules were enriched by six cycles of PCR with the following reaction composition: 2.5 U of uracil-insensitive *PfuTurboCx* Hotstart DNA polymerase (Agilent), 5 µl 10X *PfuTurbo* reaction buffer, 25 µM dNTPs, 1 µl PE Primer 1.0 (Illumina), 1 µl PE Primer 2.0 (Illumina) (50 µl final volume). The thermocycling parameters were: 95 °C 2 min, 98 °C 30 sec, then 6 cycles of 98 °C 15 sec, 60 °C 30 sec and 72 °C 4 min, ending with one 72 °C 10 min step. The reaction products were purified using the QIAquick PCR spin column (Qiagen). Two separate PCR reactions were performed on subsets of the adapter-ligated, bisulfite-converted DNA, yielding two independent libraries from the same biological sample. The quality of the libraries was checked on a Bioanalyzer followed by quantification of the libraries by qPCR using the KAPA Library Quantification Kit prior to sequencing. Samples were sequenced on an Illumina HiSeq 2000 using 50- and 59-bp single-end reads. We obtained the final sequence coverage by sequencing the two libraries for a sample separately, thus reducing the proportion of apparent PCR duplicates. The sodium bisulfite non-conversion rate was calculated as the percentage of cytosines sequenced at cytosine reference positions in the Lambda genome.

**Tet-assisted bisulfite sequencing (TAB-Seq).** TAB-Seq libraries were performed as previously described ^18^ and a detailed protocol will be provided upon request. Briefly, genomic DNA was spiked with 0.25% of *M. SsI* methylated lambda DNA and 0.25% of 5hmC spike-in control II (where all cytosines were 5hmC) and then sonicated to 200-500bp with a Covaris ultrasonicator. The *M. SsI* methylated lambda DNA and the 5hmC spike-in control were used to evaluate the conversion rate of 5mC and protection rate of 5hmC, respectively (see TAB-Seq data processing section). Next, we use b-glucosyltransferase (bGT) to introduce a glucose onto 5hmC, generating b-glucosyl-5-hydroxymethylcytosine (5gmC) to protect 5hmC from further TET oxidation. After blocking of 5hmC, all 5mC is converted to 5caC by oxidation with an excess of recombinant Tet1 protein. Bisulfite treatment of the resulting DNA then converts all C and 5caC (derived from 5mC) to uracil or 5caU, respectively, whereas the original 5hmC bases remain protected as 5gmC. Thus, following sequencing, only the 5hmC that were protected from bisulfite conversion will be read as cytosine bases. After the treatment, we performed bisulfite conversion and library preparation following a protocol identical to that for the MethylC-Seq libraries (described above). Samples were sequenced on an Illumina HiSeq 2000 using 100-bp paired-end reads.

**5hmC staining.** The protocol was adapted from Santos et al.^50^. Briefly DCs were cultured on poly-L-lysine-coated coverslips. Cells were fixed for 30 min in 4% paraformaldehyde in PBS and permeabilized with 0.2% Triton X-100 in PBS for 30 min at room temperature (RT). Cells were then washed with 0.05% Tween 20 in PBS and were treated with 1M HCl plus 0.1% Triton X-100. After 30 min at 37°C, cells were incubated with 100mM tris/HCl (pH 8.5) for 30 min and blocked for 2h in PBS with 1% BSA, 0.05% Tween-20 and 2% goat serum. Cells were incubated with 5-Hydroxymethylcytosine antibody (ActiveMotif), followed by Alexa 488 goat anti-rabbit antibody (Life Technologies) 1h at RT. The slides were mounted with Fluoromount G (SouthernBiotech), and cells counterstained with DAPI to localize the nucleus. A laser-scanning microscope (Zeiss LSM 700) in the tile scan mode was used to capture a mosaic of images. Fluorescence was quantified using the Fiji software. Average fluorescence estimates were calculated from 1,769 non-infected cells and 1,532 MTB-infected cells.

**RNA-Seq library preparation and sequencing.** RNA-Seq libraries for the six samples for which we collected MethylC-Seq were generated via polyA+ selection of mRNA from total RNA using the TruSeq RNA Sample Prep Kit v2 (Illumina). In addition, for the two individuals from whom we collected histone mark ChIP-Seq data, we also performed RNA-Seq on the whole transcriptome following ribosomal depletion using the Ribo-Zero Gold depletion and the Illumina Total Stranded RNA Library kits (Illumina). We did so in order to be able to capture enhancer RNAs, which are usually non-polyadenylated^51^. RNA-Seq libraries were sequenced as 50-bp single-end (polyA+ fraction) and 100-bp paired-end reads (ribo-minus) on an Illumina HiSeq 2500.

**ChIP-Seq library preparation and sequencing.** Samples from infected and non-infected DCs from two individuals were crosslinked with 1% w/v formaldehyde for 10 min at RT and immediately quenched for 5 min with 125mM Glycine at RT. The formaldehyde fixed samples were then sonicated to 100-400 bp using a Bioruptor (Diagenode) and then ChIP-DNA prepared using the IP-Star Compact (Diagenode) Indirect method with an Antibody-Antigen incubation of 10 hr, Bead incubation of 2 hr, and 4x 20 min wash steps. Approximately 1 million cells were used for each ChIP and ~50,000 cells for the input. The following antibodies were used: H3K4me1 (Company: CST, Cat. No.: 5326P, Lot No.: 1), H3K4me3 (CST, 9751BC, 7), H3K9me3 (MABI, 0318, 13001), H3K27me3 (MABI, 0323, 13001), H3K27ac (Abcam, Ab4729, GR119051), and H3K36me3 (MABI, 0333, 12003). ChIP and Input libraries were prepared using the Illumina Truseq Nano DNA kit, with alterations including: PCR enrichment (14 cycles) prior to size selection and utilizing the PippinPrep method (SAGE Science) instead of the SPRI method for size selection (200-400 bp). Libraries were sequenced on an Illumina Hiseq 2000. We pooled 8 libraries per lane and sequenced the lane twice to reduce the possibility of lane effects. Each library was sequenced using 50-bp single-end reads.

**ATAC-Seq library preparation and sequencing.** To prepare nuclei, we spun 100,000 cells at 500g for 5 min, which was followed by a wash using 50 µL of cold 1× PBS and centrifugation at 500g for 5 min. Cells were lysed using cold lysis buffer (10 mM Tris-HCl, pH 7.4, 10 mM NaCl, 3 mM MgCl2 and 0.05% IGEPAL CA-630). Immediately after lysis, nuclei were spun at 500g for 10 min using a refrigerated centrifuge. Immediately following the nuclei prep, the pellet was resuspended in the transposase reaction mix (25 µL 2× TD buffer, 2.5 µL transposase (Illumina) and 22.5 µL nuclease-free water). The transposition reaction was carried out for 30 min at 37 °C. Directly following transposition the sample was purified using a Qiagen MinElute kit. Following purification, we amplified library fragments using 25 uL of 2X NEBnext PCR master mix, 0.3 uL of 100X SYBR Green I, 2.5 uL each of Nextera primer index 1 (i7) and 2 (i5), and 10 uL of the transposed DNA, in a final volume of 50 uL. We used the following PCR conditions: 72 °C for 5 min; 98 °C for 30 s; and thermocycling at 98 °C for 10 s, 63 °C for 30 s and 72 °C for 1 min. To reduce GC and size bias in our PCR, we monitored the PCR reaction using qPCR in order to stop amplification before saturation. To do so, we amplified the full libraries for four cycles, after which we took an aliquot (5 ul) of the PCR reaction and added 10 µl of the previous PCR cocktail. We ran this reaction for 19 cycles to determine the additional number of cycles needed for the remaining 45 uL reaction. Libraries were amplified for a total of 9 cycles. The libraries were purified using a Qiagen MinElute kit in 20 µL. Quality of the libraries was verified on a polyacrylamide gel and Bioanalyzer. Libraries were then quantifed by qPCR using the KAPA Library Quantification Kit and sequenced on the Illumina Hiseq 2500 using 100-bp paired-end reads.

**MethylC-Seq data processing.** We used the tool Trim Galore (http://www.bioinformatics.babraham.ac.uk/projects/trim_galore/) to trim off adapter sequences incorporated in the read and remove bases with a Phred base quality score below 20. The resulting reads were mapped to the human reference genome (GRCh37/hg19) and lambda phage genome using Bismark^52^ (with the options -p 12 -N 1), which uses Bowtie 2^53^ and a bisulfite converted reference genome (C-to-T and a G-to-A) for read mapping. Only reads that had a unique alignment and a maximum number of one mismatch were retained. For each sample, we sequenced two independent libraries and therefore we removed PCR duplicates for each library separately, using a Perl script that is part of the Bismark package (deduplicate_bismark_alignment_output.pl), and then merged the two libraries for the same sample. The context of each C was determined, which allowed us to classify each C of the genome as CpG, CHH, or CHG, where H is either an A, T, or C nucleotide. Methylation levels for each CpG site were estimated by counting the number of reported C (‘methylated’ reads) divided by the total number of reported C and T (‘unmethylated’ reads) at the same position of the reference genome using Bismark’s methylation extractor tool. The same strategy was also applied for non-CpG methylation (CHG context, where H is either an A, T, or C nucleotide). We performed a strand-independent analysis of CpG methylation where counts from the two Cs in a CpG and its reverse complement (position *i* on plus strand and position *i*+1 on minus strand) were combined and assigned to the position of the C in the plus strand.

To assess MethylC-Seq bisulfite conversion rate, the frequency of unconverted cytosines (C basecalls) at lambda phage CpG reference positions was calculated from reads uniquely mapped to the lambda phage reference genome. Overall, bisulfite conversion rate was >99% in all of the samples (Table S1).

**TAB-Seq data processing.** We used Trim Galore in paired-end mode to remove adapter sequences and low quality score bases (Phred score < 20). The resulting reads were mapped in bisulfite mode to human reference genome (GRCh37/hg19) (and lambda phage + control II sequence) using Bismark with the following parameters: --bowtie2 -p 12 -N 1. PCR duplicates were removed using the deduplicate_bismark_alignment_output.pl script. In total, we obtained ~430 million paired-end reads, of which 87% were unambiguously mapped to the reference genome with a mean sequencing coverage of 10.1X and 9.3X in non-infected and infected DCs, respectively (Table S1). Similar to MethylC-Seq data, hydroxymethylation levels for each CpG site were estimated by counting the number of reported C (‘hydroxymethylated’ reads) divided by the total number of reported C and T (‘non-hydroxymethylated’ reads) at the same position of the reference genome using Bismark methylation extractor with parameters --ignore_r2 2 --no_overlap. Cytosine non-conversion rate (i.e., failed 5mC conversion by Tet1 and failure of bisulfite conversion) was assessed by calculating the frequency of C base calls at lambda CpG reference positions from reads uniquely mapped to the lambda reference. 5hmC protection rate was calculated likewise using CpG reference positions in control II sequence.

**ChIP-Seq data processing.** We started by trimming adapter sequences and low quality score bases using Trim Galore. The resulting reads were mapped to the human reference genome (GRCh37/hg19) and PCR duplicates were removed using Picard tools (http://broadinstitute.github.io/picard/). The alignment software Bowtie 2 was then used with the following options: -p 12 -N 1. Only reads that had a unique alignment and no more than one mismatch were retained. For each of the histone marks in each of the conditions, we obtained an average of 58.5 ± 9.5 SD million reads (Table S1) when combining data from the two biological replicates. Pearson correlation revealed a high concordance between the histone ChIP-Seq signals for the two biological replicates sequenced for each of the histone marks (mean r = 0.94 and range = 0.87-0.99; **Supplementary Fig. 11**), which allowed us to merge them for downstream analyses.

**RNA-Seq data processing and identification of differently expressed genes upon MTB infection.** Adaptor sequences and low quality score bases were first trimmed using Trim Galore. The resulting reads were aligned to the human genome reference sequence (GRCh37/hg19) using the TopHat2 software package^54^ with a TopHat transcript index from RefSeq. The number of read fragments overlapping with annotated exons of genes was tabulated using HTSeq^55^ using the following parameters: -q -m intersection-nonempty -s no. Using normalized gene counts for 6 infected and 6 non-infected samples, we identified genes whose expression levels were significantly altered following MTB infection of DCs using the R package DESeq2^56^. Using a paired design, we considered a gene as differentially expressed if statistically supported at a Benjamini and Hochberg^57^ false discovery rate (FDR) < 1 × 10^−4^ and showing a |log2 fold change| > 1. Lowly expressed or non-expressed genes with a read count of 0 in at least half of the samples in each condition were discarded.

**Genomic annotation and mRNA TSS collection.** Gene locations used in **Figure 1** were defined based on the GRCh37/hg19 assembly. Annotation of known Ensembl transcripts was obtained from UCSC (http://hgdownload.cse.ucsc.edu/goldenPath/hg19/database/ensGene.txt.gz). Since genes can have multiple transcripts, we selected the 5’-most transcription start site (TSS) on the positive strand as the single TSS associated with each gene. The reverse (3’ most TSS) was done for genes on the negative strand. We limited downstream analysis to protein-coding genes, resulting in 20,745 TSSs in total. Similarly, annotations for retro-elements (i.e., LINEs and SINEs), CpG islands, exons and introns were downloaded from the UCSC.

**Integrated analysis of gene expression and 5mC.** FPKM (fragments per kilobase of exon per million fragments) values of expression were calculated using Cufflinks^58^. Genes were then classified into quartiles based on their basal gene expression levels: 1st quartile is lowest and 4th is highest. Gene bodies and 20-kb regions upstream and downstream were each divided into 50 intervals. We gathered methylation data from windows within each of these intervals and plotted the mean methylation level (mean_me) for all windows overlapping each position. For each bin containing n sites (*i*):

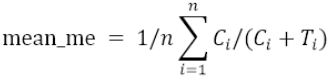

where C = read supporting methylated cytosine, T = read supporting unmethylated cytosine, *i* = position of cytosine and n = total number of cytosine positions.

**Principal component analysis based on methylomes.** Principal component analysis (PCA) of DC methylomes and those of other cell types for which MethylC-Seq data was publically available was performed on a set of 2,724,731 CpG sites that were sequenced at coverage ≥5 across all cell types or tissues using MethylKit tools^59^. The following cell types or tissues were used: neuroectoderm, neuroepithelial, glia, fetal (fheart, fthymus, fmuscle, fadrenal, fbrain), adipocyte, colon mucosa (cmucosa), substantia nigra (snigra), B-cell, T-cells (cd4, cd8, cd34), dendritic cells (dc81, dc82, dc83, dc87, dc89, dc91, hippocampus, hspc, liver, neutrophil, peripheral blood mononuclear cell (pbmc), and sperm.

**Identification of MTB-DMRs.** The summarized methylation estimates of strand-merged CpG sites from the 6 infected and 6 non-infecred samples were used to identify MTB-induced differences in methylation, using the R package Bsmooth^16^ with following parameters: ns = 25 and h = 200. Bsmooth implements a smoothing method that uses a local likelihood approach to estimate the smoothed probability of methylation at each site, taking into account the spatial correlation between nearby sites and placing greater weight on sites with higher coverage. To minimize noise in methylation estimates due to low-coverage data, we restricted the differential methylation analysis to CpG sites with coverage of ≥4 sequence reads in at least half of the DC samples in each condition, which still allowed us to interrogate changes in methylation levels at ~20 million CpG sites. Moreover, to eliminate effects caused by polymorphisms, C nucleotides that overlapped with known SNPs (dbSNP132; http://www.ncbi.nlm.nih.gov/SNP/) were removed. We identified MTB-induced differentially methylated regions (MTB-DMRs) as regions containing at least 3 consecutive CpG sites that were significantly differentially methylated using a paired t-test (|t-statistic| > 4.032 at *P* = 0.01) and exhibited at least a 10% difference in mean methylation levels between treated and untreated samples.

**Assigning MTB-DMRs to genes.** To assign each MTB-DMR to a gene, we use the following rationale: if an MTB-DMR was located within a gene body the MTB-DMR was assigned to that gene; otherwise, we assigned each MTB-DMR to the closest TSS from the center position of the MTB-DMR. If the closest TSS was further away than 250 kb the gene assigned to that MTB-DMR was not included in any of the downstream analysis.

**5hmC analysis.** Metagene profiles of 5hmC were plotted as described above for the 5mC data. To plot 5hmC profiles around MTB-DMRs, the weighted mean methylation was calculated for each contiguous 100-bp bin from 3 kb upstream to 3 kb downstream of the central position of the MTB-DMR. Only CpG sites with sequencing coverage ≥4 were included in the analyses.

**Chromatin state annotation and dynamics.** We used ChromHMM^23^ with default parameters to segment the genome into different chromatin states based on six histone modifications and ChIP input. A model was learned separately for both conditions (i.e., infected and non-infected samples), producing segmentations based on the most likely state assignment of the model. We selected a 12-state model in order to allow sufficient resolution to resolve biologically meaningful chromatin patterns. We further combined segments that had comparable histone patterns, resulting in 7 biologically meaningful chromatin states (**Supplementary Fig. 3**). To evaluate the enrichment of each chromatin state at MTB-DMRs, we first assigned each MTB-DMR to a particular chromatin state based on the chromHMM segment overlapping with its midpoint. We then calculated the frequency of MTB-DMRs that were assigned to a particular chromatin state, and normalized this value against the expected frequency based on the amount of genome covered by that state. We note that we have also performed similar analyses using a unified model that learns and defines chromatin states in both infected and non-infected DCs at the same time (in contrast to doing it separately in each condition) and all our results and conclusions remain virtually the same (**Supplementary Fig. 12**).

To test the hypothesis that regions that changed DNA methylation are also more likely to change chromatin state (compared to other regions of the genome), we randomly sampled an equal number of regions matched for the same chromatin states observed within hyper- and hypomethylated MTB-DMRs in non-infected DCs. We then counted the proportion of these random regions that changed chromatin state after infection. The expected distribution of chromHMM state transitions was generated using 1000 simulations and was compared to the proportion of chromatin changes observed among hyper- and hypomethylated regions. A similar resampling strategy was used to test for an enrichment of hypomethylated regions marked as heterochromatin/repressed before infection and that gained *de novo* enhancer marks upon MTB infection.

**Enhancer classification of hypomethylated regions based on chromatin state.** In order to define different categories of enhancers, we centered our analysis on H3K4me1 signals. If H3K4me1 was present in the basal state, such region was defined as a predefined enhancer. Therefore, predefined enhancers were simply defined as regions that overlapped with a chromHMM segment of either state 3, 4, or 5 (active or inactive enhancers) prior to MTB infection. If H3K4me1 was not found to be enriched against input in the basal (untreated) state but H3K4me1 and/or H3K27ac were induced by MTB infection, the region was defined as a *de novo* enhancer. Therefore, *de novo* enhancers were defined as regions that overlapped with a chromHMM segment of state 7 (heterochromatin/repressed) that transitioned to either state 3, 4, or 5 (active or inactive enhancers) after MTB infection.

**ChIP-Seq profiles around MTB-DMRs.** Global visualization for chromatin modifications, genome accessibility and RNA patterns around MTB-DMRs was accomplished with ngs.plot^60^ using default parameters. For each MTB-DMR, data was analyzed from 3 kb upstream to 3 kb downstream of the central position of the MTB-DMR unless otherwise indicated. To compensate for differences in total sequencing read depth among samples, all ChIP-Seq read counts were first normalized to their equivalent total number of reads. Next, the normalized number of reads was subtracted from the normalized number of reads in the input within a 100-bp scanning window, and the subtracted value was used for further analysis and plotting. For visualization purposes, pseudo counts were added if the resulting values were negative.

**Sequencing read data visualization.** Sequencing experiments were visualized by preparing custom tracks for the WashU Epigenome browser (http://epigenomegateway.wustl.edu/).

**ATAC-Seq data processing and footprinting analysis.** ATAC-Seq^29^ reads were mapped to the human reference genome (GRCh37/hg19) using BWA-MEM^61^, at default parameters. Only reads that had a unique alignment and mapping quality of ≥10 were retained. Similarly, ngs.plot was used to plot ATAC-Seq profiles around MTB-DMRs. To detect TF binding footprints in the ATAC-Seq data we used the program Centipede^31^ in two steps. In the first step, we determined which transcription factors were active (have motif instances with footprints) before and after infection using a reduced set of motif instances (5K-15K) for each TF as defined in Moyerbrailean *et al*.^62^. In the second step, we scanned the entire genome for motif instances matching the original PWM, and we ran Centipede in parallel for the two conditions in order to make the posterior probabilities comparable. For both steps, to run Centipede the aligned paired-end reads are separated into four bins depending on the fragment length ([40, 99], [100, 139], [140, 179] and [180, 250] in bp). As Tn5 transposase contacts and duplicates 9 bp of DNA^29^ we take as the cleavage site the middle nucleotide. To do so, we shifted 4 bp from the 5’-end positions towards the center of the fragment. Then for each motif we built a matrix that counted Tn5 cleavage events, where each row represented a motif instance (i.e., a candidate binding site), and each column represented a spatial location with respect to the TF binding site in bp (i.e., relative cleavage site). This matrix was constructed separately for each fragment length bin and each strand orientation (with respect to the motif match, or reference strand if motif was palindromic). We used a window size of 300 bp on either side of the motif match. We then concatenated all 8 matrices and fed them as input data to Centipede, together with the PWM score.

For determining which TFs are active on the first step, we calculate a Z-score corresponding to the PWM effect in the prior probability in Centipede’s logistic model and we determine as active those that have a Bonferroni-corrected *P* < 0.05. The Z-score corresponds to the *β* parameter in:

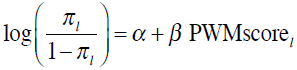

where *π*_*l*_ represent the prior probability of binding in Centipede’s model in motif location *l*. On the second step, we first train Centipede assuming that the footprint is bound in the two conditions. Then, we fix the model parameters and we generate a likelihood ratio and posterior probability *π*_*lt*_ for each condition *t* separately and for each site *l*. To detect if the footprint is more active in one of the two conditions, we fit a logistic model that includes an intercept for each condition (*α* and *β*), the PWM effect *β*, and PWM times the treatment effect *γ*:

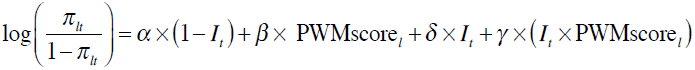

where *I*_*t*_ is an indicator variable that takes the value 1 if t = “treatment” and 0 if t = “control’’. We calculate a Z-score for the interaction effect *γ* that measures if there is condition specific binding.

**Gene Set Enrichment Analysis.** We used Genetrail^63^ to test for enrichment of functional annotations among genes nearby MTB-DMRs (<250 kb), using the set of all Ensembl genes as a background. Analysis was done with default parameters and results corrected for multiple testing by the method of Benjamini and Hochberg to control the False Discovery Rate (FDR).

